# Non-coding RNA Repertoire in Reef-Building Corals

**DOI:** 10.1101/2025.03.15.643469

**Authors:** Jill Ashey, Javier A. Rodriguez-Casariego, Kathleen M. Durkin, Zachary Bengtsson, Samuel J. White, Ariana S. Huffmyer, Danielle M. Becker, Jose M. Eirin-Lopez, Hollie M. Putnam, Steven B. Roberts

**Affiliations:** Department of Biological Sciences, University of Rhode Island, Kingston RI USA; Environmental Epigenetics Laboratory, Institute of Environment, Florida International University, Miami FL USA; Rosenstiel School of Marine, Atmospheric and Earth Sciences. University of Miami, Miami FL USA; School of Aquatic and Fisheries Sciences, University of Washington, Seattle WA USA

**Keywords:** non-coding RNAs, long ncRNAs, microRNAs, piwiRNAs, corals

## Abstract

Non-coding RNAs (ncRNAs) play critical regulatory roles in gene expression regulation that influences diverse biological processes in response to environmental change. Yet their characterization in non-model organisms, particularly sessile, benthic ecosystem engineers such as reef-building corals that are sensitive to climate change, remains limited. This study provides the first comprehensive analysis of the ncRNA repertoire of species from three ecologically important coral genera from Mo’orea, French Polynesia: *Acropora pulchra*, *Pocillopora tuahiniensis*, and *Porites evermanni*. These species demonstrate differing symbiotic partners, life history strategies, and physiological traits, offering a broad framework for documenting ncRNA variation in corals. We identified homologs for ncRNA biogenesis and functional machinery, characterized long ncRNAs (lncRNAs), microRNAs (miRNAs), and piwiRNAs (piRNAs), and assessed their genomic context and potential targets. Our findings reveal the presence of conserved ncRNA machinery across these coral species, indicating their capability to generate and utilize ncRNAs for the regulation of gene expression. We identified only a single miRNA conserved with corals and Eumetazoans (miR-100), four miRNAs shared across all three species, previously identified in other cnidarian taxa (miR-100, miR-2023, miR-2025, miR-2036), as well as several species-specific miRNAs. Predicted gene targets of the characterized miRNAs included immune response regulation in *A. pulchra* and *P. tuahiniensis* and signal transduction pathways in *P. evermanni* and *P. tuahiniensis*. Proximity analysis indicated >71-99% of piRNAs overlapped with genes, with genomic maintenance and stability identified as the primary functional enrichment of those genes. Our characterization of lncRNAs found little sequence overlap across each species (<2%), although lncRNAs in all three species were often in proximity to immune-related genes. This study lays the groundwork for the repertoire and regulatory roles of ncRNAs in reef-building corals, thereby expanding our understanding of epigenetic regulation in environmentally sensitive marine invertebrates and its potential implications in acclimatization and adaptation to environmental change.

## Introduction

Long thought to be the molecular “junk” of the cell, non-coding RNAs have emerged as pivotal regulators of cellular structure and function, via key roles in gene expression regulation across diverse cell types and tissues (Eddy, 2002). Unlike protein-coding RNAs (i.e., mRNAs), non-coding RNAs do not encode for proteins, but instead perform crucial structural and regulatory functions (Hüttenhofer et al., 2005). Non-coding RNAs can be categorized into two broad classes: structural RNAs, such as ribosomal RNAs (rRNAs) and transfer RNAs (tRNAs) that primarily facilitate translation through their secondary structures, and regulatory ncRNAs, including long and small non-coding RNAs (ncRNAs) (Hüttenhofer et al., 2005) that interact with mRNA transcripts to regulate their translation or degradation. Regulatory ncRNAs can be further subdivided into different families based on their functions in the cell and the length of the transcript, typically being categorized as long non-coding RNA (lncRNA; >200 nucleotides) and small non-coding RNA (sncRNA; <200 nucleotides). sncRNAs can be further divided into microRNAs (miRNAs, ∼22 nucleotides) that regulate mRNA translation and degradation, piwi interacting RNAs (piRNAs, ∼24-32 nucleotides) with a silencing role, and other nuclear ncRNAs, such as small nuclear RNAs (snRNAs, ∼150 nucleotides), which aid in spliceosome assembly and function in the nucleus (Bohnsack & Sloan, 2018; Gomes et al., 2013).

Regulatory ncRNAs interact with mRNA via direct and indirect mechanisms. miRNAs interact with Argonaute proteins (i.e., family of proteins involved in gene expression regulation) to form the RNA-induced silencing complex (RISC), which, upon binding to the mRNA 3’ untranslated region (3’UTR), facilitates translational repression or degradation (Kaikkonen et al., 2011). Similarly, piRNAs associate with Piwi-proteins (Argonaute/PIWI family) to form piRISC complexes that silence transposable elements and other complementary RNA targets (e.g. pseudogenes, *nanos* mRNA, transcripts) in the germline and the soma (Czech et al., 2018; Czech & Hannon, 2016). lncRNA function is broader in scope than other ncRNAs, with evidence that they play key regulatory roles in gene expression, chromatin organization, and cellular architecture, through RNA-RNA, RNA-DNA, and RNA-protein interactions (Mattick et al., 2023).

In plants and animals, ncRNAs play a critical role in gene expression regulation in response to environmental perturbations (Chao et al., 2022; Fu, 2014; Gomes et al., 2013). For example, depletion of a miRNA (miR-85) in *Caenorhabditis elegans* during heat stress resulted in overexpression of heat shock proteins and reduced survival (Pagliuso et al., 2021). In *Arabidopsis*, the knockdown of a lncRNA (COLD INDUCED lncRNA 1) increased susceptibility to cold stress, as evidenced by increased endogenous reactive oxygen species and reduced osmoregulatory substances (Liu et al. 2022). Further, piRNAs represses expression of transposable elements in *Hydra* somatic cells, potentially contributing to their longevity of *Hydra* (Teefy et al., 2020). These examples across a range of phyla and kingdoms demonstrate that ncRNAs play important roles in phenotypic responses to environmental change, and thus ncRNA presence and regulatory capacity is likely to have a role in environmental response in reef-building corals.

While studies in model organisms have built the basis of our knowledge of ncRNAs, non-model organisms such as aquatic/marine, sessile, benthic taxa that require acclimatory mechanisms for persistence have been less explored (Leitão et al., 2020). Reef-building corals, in the phylum Cnidaria, are key ecosystem engineers of massive ecosystems, are basal metazoans, and are sessile for the majority of their lives. Therefore, they must modulate their physiological state in response to seasonal and diel natural variability (Barshis et al., 2013; Brown et al., 2022; Safaie et al., 2018), as well as to stochastic environmental changes due to both natural disturbances and increasing anthropogenic stressors (Ellis et al., 2019; Grigg & Dollar, 1994; Richmond et al., 2018; Sweet & Brown, 2016). Further, reef-building corals are highly susceptible to increasing sea surface temperatures and heatwaves (Gutierrez et al., 2024; Mellin et al., 2024), which causes coral bleaching or a breakdown in the coral and endosymbiotic dinoflagellate symbiosis (Gates et al., 1992) that provides the energy for coral growth and reproduction (Cunning et al., 2017; Edmunds & Davies, 1986; Osinga et al., 2011).

Given the central role of ncRNAs in gene expression regulation in other species, the identification and characterization of ncRNAs in corals will provide a more clear understanding of the regulatory mechanisms governing coral gene expression regulation and stress response, and further provide insight into the evolution of RNA-mediated regulatory networks across metazoans. To date, there has been one study focused on the miRNA repertoire in a single species of coral (Liew et al., 2014), which found potential roles for miRNAs in the regulation of symbiosis and calcification. Further, both miRNAs and lncRNAs have been implicated in thermal stress responses in single coral species studies (Gajigan & Conaco, 2017; Huang et al., 2017, 2019; Yu et al., 2021). Unraveling the roles of ncRNAs in coral stress responses is essential for assessing their contributions to coral resilience and, more broadly, to the stability of reef ecosystems facing environmental change.

Here, we provide the first in-depth characterization of ncRNAs in three important reef-building coral genera: *Acropora*, *Pocillopora*, and *Porites*. We chose three species (*Acropora pulchra*, *Pocillopora tuahiniensis*, and *Porites evermanni*) that are representative of genera commonly found on reefs worldwide and are evolutionary distant (Kitahara et al., 2010; Stolarski et al., 2011). Further, these species display contrasting symbiotic and life history strategies and thermal sensitivities, presenting a compelling system to examine variation in ncRNA in corals. For instance, branching *Pocillopora* (robust clade; (Kitahara et al., 2010)) and *Acropora* spp. (complex clade; (Kitahara et al., 2010)) are generally more sensitive to environmental changes than massive *Porites* spp. (complex clade; (Kitahara et al., 2010)), which tend to exhibit greater environmental tolerance (Loya et al., 2001). These species also differ in their association with Symbiodiniaceae taxa with *Pocillopora* and *Acropora* spp. as symbiotic generalists hosting a mixture of *Cladocopium*, *Symbiodinium*, and/or *Durusdinium* while *Porites* exhibits high fidelity with specific *Cladocopium* symbionts (e.g., C15) (Putnam et al., 2012). Thus, the similarity or divergence of coral ncRNA repertoires could indicate their involvement in modulating such phenotypic traits, and its relevance to other coral genera.

In this study, we aimed to:

1. Identify homologs for ncRNA-related protein machinery to understand the types of ncRNAs present and their potential functions.
2. Characterize the features of ncRNAs, focusing on the lncRNAs, miRNAs, and piRNAs, and assess sequence similarities between species.
3. Identify the nearest gene to lncRNAs and piRNAs and explore potential functional roles of ncRNA regulation.
4. Determine putative gene targets of miRNAs to elucidate the potential biological processes and pathways affected by miRNA-mediated regulation.

## Methods

### Study site and collection

Sample fragments of three coral species (*Acropora pulchra*, *Pocillopora tuahiniensis*, and *Porites evermanni*) were collected from the lagoon backreef in Mo’orea, French Polynesia on March 5, 2020 on the north shore (−17.476872, −149.80594). These species are ecologically common on French Polynesian reefs, with great importance as habitat builders (Vercelloni et al., 2019). During sampling, n=1 fragment (1 cm) was collected from five colonies of each species using bone clippers, snap frozen in liquid nitrogen and then put into 1.5 mL tubes with 600 μL of Zymo DNA/RNA Shield (Zymo CAT R1100-250) and stored at −40°C at the University of California Berkeley Richard B. Gump South Pacific Research Station until transportation to the University of Rhode Island for further processing. Samples were exported under CITES FR2198700194-E on 04/06/2022.

### Species Identification

Cryptic species are common in reef-building corals (Burgess et al., 2021, 2024; Forsman et al., 2009). *Acropora pulchra* has historically reliably been identified by morphology and was transplanted to the reef site from a coral nursery, so we sequenced a single sample for validation. One *A. pulchra* colony sample was sequenced using Sanger sequencing by amplifying the *Pax*-C 46/47 intron in the nuclear genome region as described by (Van Oppen et al., 2001, 2004) as well as the mitochondrial putative control region (933+ bp) plus 83 bp of cytochrome oxidase III as described by (Vollmer & Palumbi, 2002). *Pocillopora spp* and *Porites spp,* are known to include cryptic species, requiring genetic identification (Burgess et al., 2021, 2024; Forsman et al., 2009). *Pocillopora* species were identified by amplifying the mitochondrial open reading frame (mtORF) region as described by (Burgess et al., 2021; Johnston et al., 2018). *Porites* species groups (*P. evermanni* vs *P. lobata*/*P. lutea*) were identified using the coral nuclear histone region spanning H2A to H4 (i.e., H2) (Tisthammer et al., 2020). Species identification methods can be found in the Supplemental Methods. Sanger sequences have been deposited at https://osf.io/aw53f/.

### Extractions & Sequencing

DNA and RNA were extracted using the Zymo Quick-DNA/RNA Miniprep Plus Kit (Zymo CAT R1057) according to the manufacturer’s instructions. DNA and RNA integrity and quantity were assessed using a 1.5% agarose gel and a Qubit fluorometer, respectively. Sequencing was performed by Azenta Life Sciences. For RNA sequencing (which identifies lncRNAs), strand-specific RNA sequencing libraries were prepared using NEBNext Ultra II Directional RNA Library Prep Kit for Illumina following the manufacturer’s instructions (NEB, Ipswich, MA, USA). Libraries were sequenced using a 2×150bp Paired End (PE) configuration on Illumina HiSeq 4000. For short RNA sequencing, sequencing libraries were prepared using NEB Small RNA Library Prep Kit (NEB CAT: E7560S) following the manufacturer’s instructions (New England BioLabs, Ipswich, MA, USA). Libraries were sequenced using a 2×150bp Paired End (PE) configuration on Illumina HiSeq 4000.

### Genomic resources

Genomes, protein and gene sequences, and annotation files were downloaded from the following sources. At the time of analysis, *Acropora pulchra* did not have a published genome, so the genomic information from *Acropora millepora* was used for the analysis. *Acropora millepora* genomic information was downloaded from NCBI (PRJNA633778) and has an estimated genome size of 475 Mbp, N50 of 19.8 Mbp, and 28,188 protein coding genes (Fuller et al., 2020). While *A. millepora* and *A. pulchra* are morphologically distinct, their close evolutionary relationship and ability to interbreed (Van Oppen et al., 2001, 2002; Wallace & Willis, 1994) support the use of the *A. millepora* genome as a reference for *A. pulchra* analyses. Currently to our knowledge, *Pocillopora tuahiniensis* does not have a published genome, so the genomic information from *Pocillopora meandrina*, a closely related species, was used for the analysis. *Pocillopora meandrina* genomic information was downloaded from http://cyanophora.rutgers.edu/Pocillopora_meandrina/ with an estimated genome size of 377 Mbp, N50 of 10 Mbp, and 31,840 protein coding genes (Stephens et al., 2022). *Pocillopora meandrina* and *Pocillopora tuahiniensis* are morphologically similar, and despite being in different clades, mitochondrial and nuclear genomic data demonstrate that they are closely related; this evolutionary relationship supports the use of the *P. meandrina* genome as a reference for *P. tuahiniensis* (Johnston & Burgess, 2023). While *P. verrucosa* is more closely related to *P. tuahiniensis* (Johnston & Burgess, 2023) and has a published genome (Buitrago-López et al., 2020), our analysis used *P. meandrina* due to its significantly higher assembly contiguity (*P. meandrina* N50 = 10 Mbp, *P. verrucosa* N50 = 0.334 Mbp) and more extensive genome annotation. These factors make the *P. meandrina* a more suitable reference genome for this study. *Porites evermanni* genomic information (generated from a sample from Mo’orea, (Noel et al., 2023)) was downloaded from https://www.genoscope.cns.fr/corals/genomes.html with an estimated genome size of 497 Mbp, N50 of 0.17 Mbp, and 40,389 protein coding genes (Noel et al., 2023). Versions of the genomic resources used in this study are archived at https://osf.io/aw53f/.

### ncRNA protein machinery identification

To validate the presence of specific ncRNAs in these coral species, homologs of ncRNA-related proteins of interest were identified in the target coral species genomes, including: ribonuclease III (Drosha), microprocessor complex subunit (DCGR8), exportin-5 (XPO5), endoribonuclease (Dicer), argonaute-2 (AGO2), piwi, phosphatidylinositol-4-phosphate 5-kinase type 1 alpha (PIP5K1A), and ribonuclease P (RNase P). The reference sequences of proteins of interest were obtained from six species (*Pocillopora verrucosa*, *Orbicella faveolata*, *Acropora millepora*, *Stylophora pistillata*, *Nematostella vectensis*, and *Homo sapiens)* available on NCBI (Table S1), and were compared against the protein fasta for each of the three target species genomes above using blastp (evalue 1E^−40^, max_target_seqs 1, max_hsps 1; (Altschul et al., 1990).

### Long ncRNA characterization

RNA sequencing read quality was initially assessed using FastQC (Andrews 2010; version 0.12.1) and MultiQC (Ewels et al., 2016). RNA sequencing reads underwent quality trimming using Fastp (Chen et al., 2018), which involved the identification of read pairs, automatic detection and removal of adapter sequences, trimming of poly-G tails, and removal of the first 20 bases from both ends to eliminate low-quality bases. Trimmed reads were aligned to respective genomes with HISAT2 (Kim et al., 2019) and assembled with Stringtie (M. Pertea et al., 2015) to obtain read counts and merge sample GTF files. Sample GTF files were then compared to genome annotations using gffcompare (G. Pertea & Pertea, 2020) and filtered so that only transcripts >199 bp were kept for analysis. Coding Potential Calculator 2 was implemented to predict whether a transcript was coding or non-coding (Kang et al., 2017), and coding transcripts were removed from analysis. In all cases, default parameters were used. Putative lncRNA GTF and fasta files were generated for each species. In order to evaluate sequence similarity across species, lncRNAs from each species were compared to the other species using blastn (evalue 1E^−40^, max_target_seqs 1, max_hsps 1; (Altschul et al., 1990); version 2.15.0).

### Short ncRNA: miRNA characterization

To characterize miRNAs, the quality of short RNA sequence reads was assessed using FastQC (Andrews 2010) and MultiQC (Ewels et al., 2016). Fastp default parameters were used for read quality trimming (Chen et al., 2018), which involved the removal of adapter sequences (provided from the NEB Small RNA Library Prep Kit) and polyG sequences. Reads with a length greater than 31 bp were discarded; Read 1 and Read 2 were then merged (using --merge flag from fastp), with 17 bp as the minimum overlap (Chen et al., 2018).

To identify previously described miRNAs, a database containing mature miRNA sequences from miRBase (https://www.mirbase.org/; version 22; (Kozomara et al., 2019)) and cnidarian miRNA sequences manually collected from published articles (Table S2) was used as a reference. Previously described and candidate novel miRNAs were identified from alignments of trimmed and filtered reads to respective reference genomes using ShortStack (v4.0.2, (Axtell, 2013; Shahid & Axtell, 2014)) with the parameter –dn_mirna, which identifies putative novel miRNAs by searching read-genome alignments for “islands” of significant coverage and evaluating neighboring regions based on a stringent set of miRNA precursor gene characteristics. To identify miRNAs conserved within all or a subset of the three study species, blastn was run with the –task blastn-short option (evalue 1E^−40^, max_target_seqs 1, max_hsps 1; Altschul et al.

1990).

### Short ncRNA: Piwi RNA characterization

For piRNA characterization, quality filtered and trimmed short reads (*“Short ncRNA: miRNA characterization”* section) were filtered to keep reads between 25 and 31 bp. Low complexity sequences were removed using NGS toolbox scripts (Rosenkranz et al., 2015). To remove non-piRNA, short reads were mapped to a track of other non coding RNAs with SortMeRNA (v4.3.6, (Kopylova et al., 2012)). Reference tRNA, rRNA, miRNA and siRNA sequences were acquired or generated for each species. For *A. pulchra*, tRNA and rRNA sequences were obtained from annotation files from the *A. millepora* genomic resources; for *P. tuahiniensis* and *P. evermanni*, tRNA and rRNA sequences were generated *de-novo* using tRNAscan-SE (v.2.0.9; Chan et al. 2021) and Barrnap (v.0.9; https://github.com/tseemann/barrnap) with default parameters for eukaryotes. Hairpin, star, and mature miRNAs, and siRNAs annotations were obtained from ShortStack output as described above. Sequences that did not align to these known non-coding RNAs were mapped to the respective genome using sRNAmappper.pl from the NGSToolbox using the option -alignments best, and origin assignment for reads with multiple mappings was determined using reallocate.pl (Rosenkranz et al., 2015; Rosenkranz & Zischler, 2012). The resulting sequences were considered putative piRNAs and used for downstream analyses.

The following analysis was used to identify putative piRNA clusters in *A. pulchra*, *P. tuahiniensis*, and *P. evermanni*. ProTrac (v2.4.4) was used to predict piRNA clusters for each sample (Rosenkranz & Zischler, 2012). Resulting piRNA clusters were merged by species if they were present in at least two replicates, with a minimum overlap of 50% of the cluster length. PPmeter (v0.4, (Jehn et al., 2018)) was used with default parameters to quantify and compare ping-pong amplification signatures, a hallmark of secondary piRNA biogenesis (Czech & Hannon, 2016). In the ping-pong cycle, piRNAs are generated through reciprocal cleavage events, which creates the characteristic 10 base sequence overlap between piRNAs (Czech & Hannon, 2016).

In eukaryotes, piRNAs typically are located in transposable element (TE) regions. To investigate if piRNA clusters were located in TE in corals, TE annotation was performed in the reference genome of each species using RepeatMasker (v4.1.0, http://repeatmasker.org) with default parameters. TE enrichment in piRNA clusters was calculated respectively to each genome by quantifying annotated TE overlap with piRNA clusters and contrasting piRNA cluster-TE overlaps with respect to TE coverage across each genome using bedtools (Quinlan & Hall, 2010).

### lncRNA and piRNA genomic proximity to genes

To investigate the proximity of ncRNAs to genes, bedtools closest (v2.30.0) was performed using the GFF files downloaded for each species and the ncRNA GFF files generated from our analyses (Quinlan & Hall, 2010). The percentage of ncRNAs that did and did not overlap with a gene were calculated. If an ncRNA did overlap with a gene, average overlap in base pairs (bp) was calculated. If an ncRNA did not overlap with a gene, average distance between the two features in bp was calculated.

Functional enrichment was performed using gene ontology (GO) enrichment analyses with TopGO (Alexa & Rahnenfuhrer 2024) on the genes that were closest to or overlapping ncRNAs. The use of ‘closest’ will herein refer to either the closest gene or the overlapping gene, depending on where the ncRNA is in the genome relative to the gene. GO enrichment was run using Fisher’s exact test (weight01 algorithm with a threshold of p<0.05) to assess GO enrichment in Biological Processes for each species. Enriched GO terms were compared across species to investigate shared enriched gene functions related to Biological Processes.

### miRNA target identification

In cnidarians, miRNAs typically require binding to the 3’ untranslated region (3’UTR) of mRNAs with almost perfect complementarity to repress or downgrade translation (Moran et al., 2014). Therefore, target prediction is less prone to false positives in this group. Given the lack of 3’UTR annotation on these genomes, 1000bp on the right flank of mRNA tracks was classified as the 3’UTR. If a 3’UTR was overlapping a contiguous gene, the overlapping portions of those 3’UTRs were subtracted using bedtools (Quinlan & Hall, 2010). 3’UTR sequences were annotated from the respective genomes using bedtools getfasta (Quinlan & Hall, 2010). MiRanda was then used to predict miRNA gene targets using the constructed 3’UTR sequences and mature miRNA sequences as input with strict binding invoked (-en −20, -sc 140; (Enright et al., 2003)). Similar to the previous GO enrichment analysis, TopGO was used to assess GO enrichment in Biological Processes on the predicted miRNA-targeted genes (Alexa & Rahnenfuhrer 2024). GO enrichment analyses were also conducted on the genes targeted by the miRNAs shared across species (mir-100, mir-2023, mir-2025, mir-2036). Enriched GO terms were compared across species to investigate shared enriched gene functions related to Biological Processes for the miRNA targets.

## Results

### Species ID

mtORF haplotype ID identified all of the Pocillopora samples as *Pocillopora tuahiniensis* (Haplotype_10; Johnston and Burgess 2023; Figure S1). Coral nuclear histone region spanning H2A to H4 (H2; (Tisthammer et al., 2020)) indicated all of the *Porites* samples group with the reference sequences for *Porites evermanni* (Figure S2).

### ncRNA protein machinery

At least one protein homolog of all ncRNA proteins of interest (Table 1) was identified in each species in this study.

**Table 1.** Presence of ncRNA protein machinery in each species. The number indicates the number of unique proteins of that type were found.

| Protein | Function | *A. pulchra* | *P. evermanni* | *P. tuahiniensis* |
| --- | --- | --- | --- | --- |
| AGO2 | Part of miRNA/siRNA silencing complex | 10 | 2 | 3 |
| DGCR8 | Forms microprocessor complex with Drosha that cleaves the pri-miRNA into a pre-miRNA (stem-loop structure) | 3 | 1 | 1 |
| Dicer | Cleaves loop from pre-miRNA to form mature miRNA | 4 | 4 | 5 |
| Drosha | Forms microprocessor complex with DGCR8 that cleaves the pri-miRNA into a pre-miRNA (stem-loop structure) | 2 | 2 | 2 |
| Pip5k1a | Blocks pre-miRNA transport into cytoplasm | 3 | 3 | 2 |
| Piwi | Interacts with piRNAs, facilitates 'ping-pong' piRNA biogenesis | 3 | 2 | 2 |
| Rnase P | Cleaves lncRNA to produce mature 3' ends | 9 | 8 | 7 |
| Xpo5 | Exports pre-miRNA from nucleus to cytoplasm | 8 | 1 | 1 |

### Long ncRNA

Following filtering for size (>199nt), as well as excluding known coding transcripts and predicted coding sequences, the number of putative lncRNA transcripts identified was 16,206 in *A. pulchra*, 7,378 in *P. evermanni*, and 12,395 in *P. tuahiniensis.* The mean lncRNA lengths for *A. pulchra*, *P. evermanni*, and *P. tuahiniensis* were 1,058 bp, 2,450 bp, and 2,755 bp, respectively, while the median lengths were 451 bp, 930 bp, and 690 bp, respectively (Figure 1A). Comparison of shared lncRNAs between the three species indicated 648 (1.8%) shared transcripts, whereas 14,715 (40.9%), 6,836 (19%), and 11,153 (31%) lncRNAs were unique to *A. pulchra*, *P. evermanni*, and *P. tuahiniensis*, respectively. *Acropora pulchra* shared fewer lncRNAs with *P. evermanni* (1,115; 3.1%) and more lncRNAs with *P. tuahiniensis* (1,151; 3.2%). *P. evermanni* and *P. tuahiniensis* shared the fewest lncRNAs (396; 1.1%).

**Figure 1.**
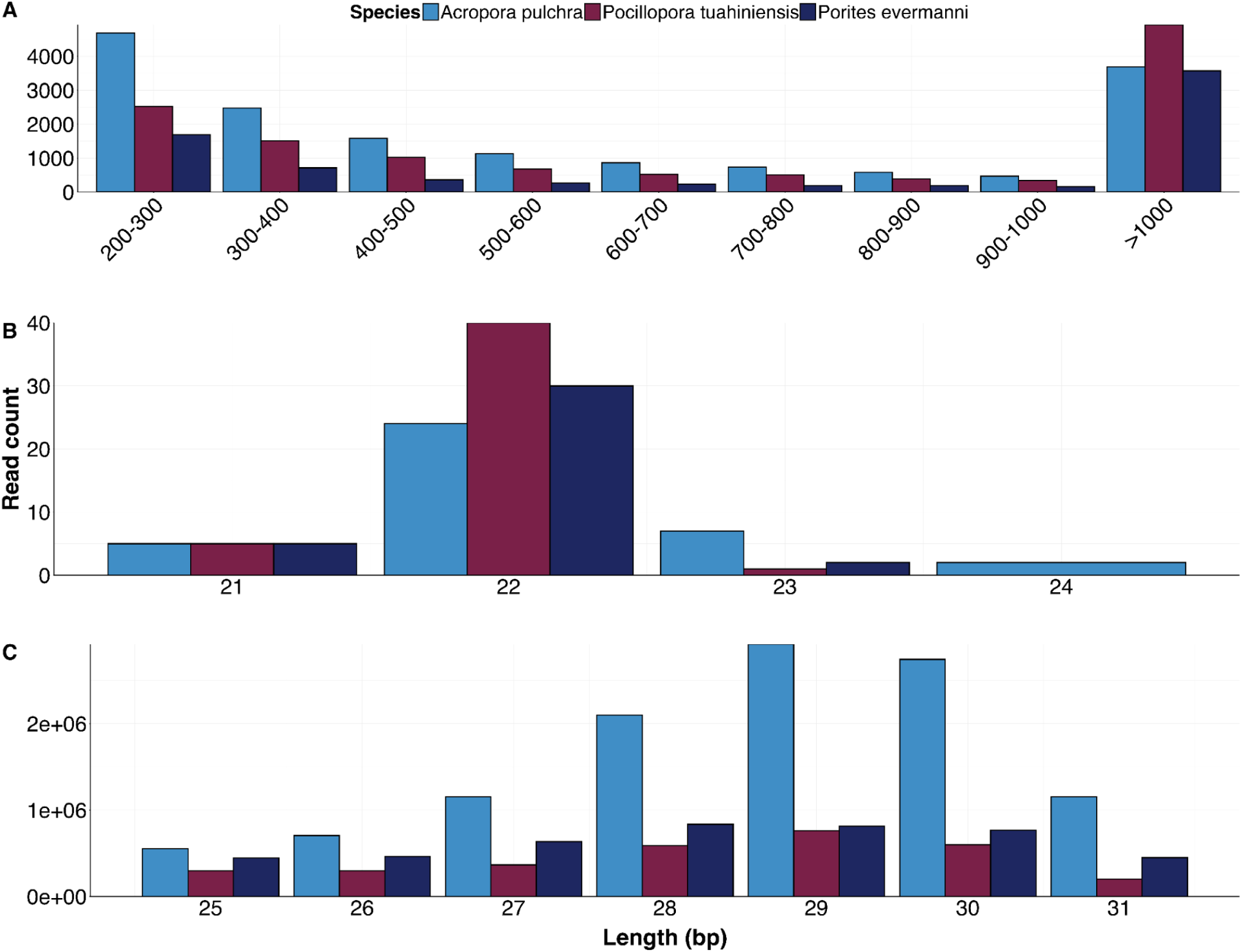
Length distribution of ncRNAs across species. A) Histogram of lncRNA lengths by species. B) Histogram of miRNA lengths by species. C) Histogram of piRNA lengths by species.

### Short ncRNA: miRNAs

A total of 38, 46, and 37 putative miRNAs were identified from *A. pulchra*, *P. evermanni*, and *P. tuahiniensis*, respectively. In all species, these miRNAs were 21 to 24 nucleotides long, with an average length of 22 nucleotides (Figure 1B). Of these miRNAs, 24 from *A. pulchra*, 9 from *P. evermanni*, and 9 from *P. tuahiniensis*, respectively, had matches to the curated database of previously described cnidarian miRNAs. The miRNAs that matched known miRNA sequences from the reference database were named after the known miRNA (i.e., if sequence matched nve-mir-100 and/or adi-mir-100, that miRNA sequence was named [coral species]-mir-100). If a sequence did not match any known sequence, it was named [coral species]-mir-novel-# in sequential order (Table S3). Sequences were identified that had identical mature sequences, but unique locations in the genome in *A. pulchra* and *P. evermanni*; these sequences were given the same number and differentiated with an “a’’ or “b”. *P. tuahiniensis* did not have any identical sequences in their putative miRNAs.

Four miRNA sequences were shared among our three species, corresponding to previously described miRNAs in Eumetazoans mir-100, and in the cnidarian *Nematostella vectensis*: mir-2023, mir-2025, and mir-2036. Additionally, *A. pulchra* and *P. evermanni* shared 1 miRNA (mir-2030); *P. evermanni* and *P. tuahiniensis* shared 1 miRNA (mir-novel-20); and *A. pulchra* and *P. tuahiniensis* shared 1 miRNA (mir-novel-7).

### Short ncRNA: piRNAs

piRNA sequence analyses identified 38.2, 10.7 and 5.4 million sequences with lengths between 24-31 bp in *A. pulchra*, *P. tuahiniensis*, and *P. evermanni*, respectively. These sequences did not match mRNA nor any other class of non-coding RNAs and were thus considered putative piRNA for further analyses. Putative piRNAs were between 24-31 bp in length, with most sequences having between 28 and 30 bp (Figure 1C). Evidence of ping-pong amplification signatures, which are indicative of secondary piRNA biogenesis, was observed in the putative piRNA transcripts through a sequence bias towards uracil on the 5’end and an adenine in the 10th position (Figure S3; (Siomi et al., 2011)).

piRNA cluster prediction was applied for reliable characterization of putative piRNAs primary transcripts. A total of 97, 107, and 47 piRNA clusters were identified in *A. pulchra*, *P. evermanni*, and *P. tuahiniensis*, respectively (Table S4). Around 30% of putative-piRNA reads were included in clusters for *A. pulchra* and *P. evermanni*, while for *P. tuahiniensis* less than 25% of the total reads were included in clusters. There was also variability in the proportion of unique sequences included in clusters per species, with *A. pulchra* having less than 20% (Fig 2A). In all cases we observed the signature sequence bias towards an uracil at the 5’ end (Fig 2A).

**Figure 2.**
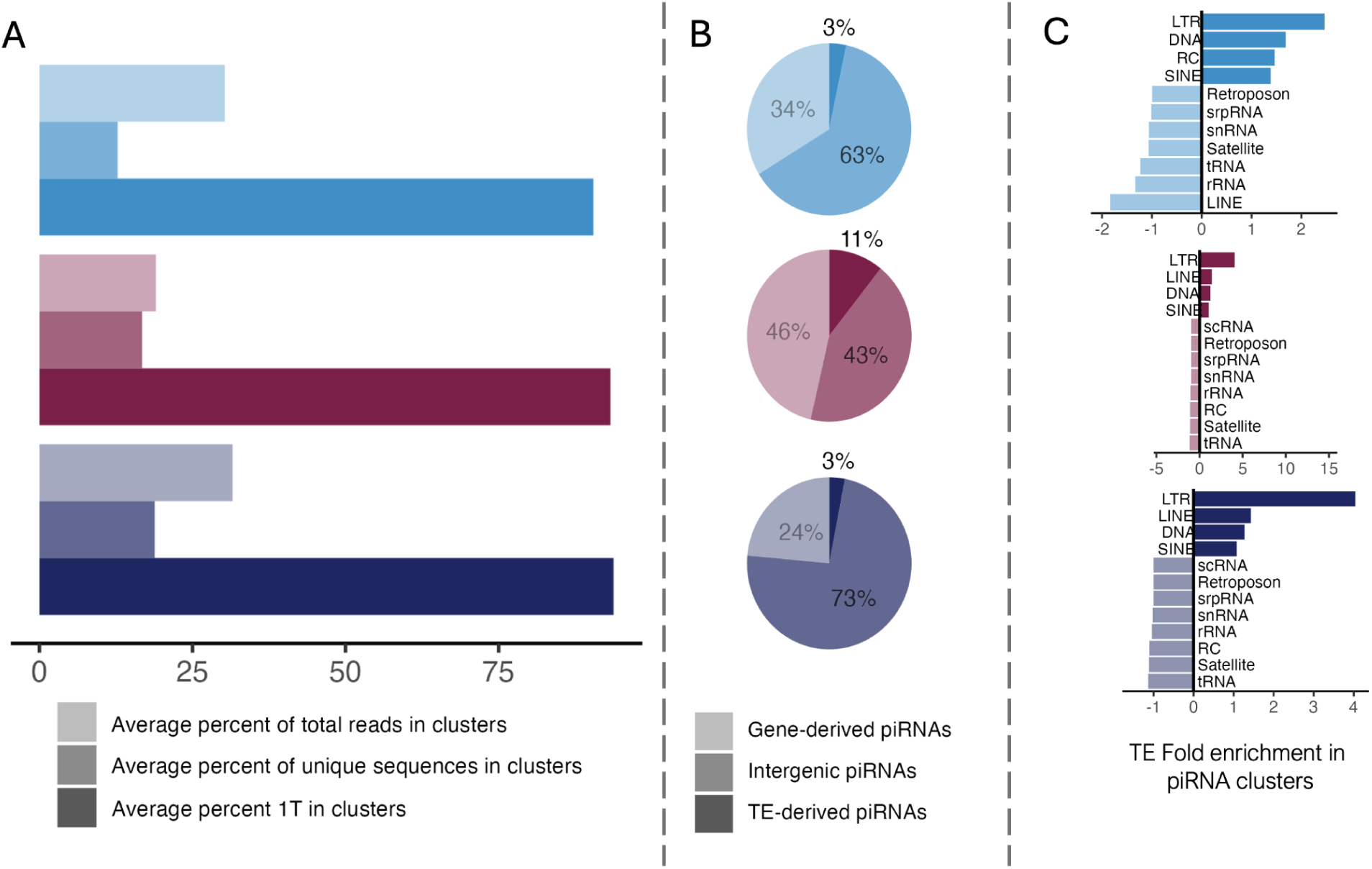
piRNA clusters characteristics in three coral species. **A.** Proportion of reads and unique sequences of putative-piRNA included in clusters, and proportion of reads in clusters with Thymine (Uracil in RNA sequence) in position 1 (5’-end) **B.** Proportions of gene-derived, repetitive element-derived, and intergenic putative piRNA sequences in clusters. **C.** Transposon enrichment in piRNA clusters respective to their abundance across the genome, indicating which families are more often targeted by piRNA regulation.

Overlaps of clusters with genomic regions of interest followed a similar pattern to all putative piRNAs (Fig 2B, Figure S4) with genes and intergenic elements being highly represented. TEs were less abundant in the piRNA clusters than in the putative-piRNA pool, except for *P. tuahiniensis* where TEs were 11% more abundant in the clusters, compared to the putative-piRNA pool. Only a small portion of transposon families were enriched in clusters respective to their presence in the genome (Figure 2C). LTR (long terminal repeats) and DNA transposons were significantly enriched in the clusters of all three species, while LINE (long interspersed nuclear element), RC (rolling-circle transposons) and SINE (short interspersed nuclear elements) transposons were differentially enriched between species.

### lncRNA and piRNA genomic proximity to genes

Across all species, 97.5% of lncRNAs did not overlap with their closest gene (Table 2). Genes closest to lncRNAs in all species were enriched for two GO biological processes: positive regulation of transcription by RNA polymerase and positive regulation of interleukin-1 beta production (Figure 3; Table S5). Additionally, genes closest to lncRNAs in *A. pulchra* and *P. tuahiniensis* showed enrichment for processes related to NF-kappaB signaling, transcription factors, and the regulation of inflammatory and innate immune responses (Figure 3; Table S5). In *A. pulchra* and *P. evermanni*, genes closest to lncRNAs were enriched for processes involved in wound healing and I-kappaB kinase/NF-kappaB signal transduction (Figure 3; Table S5).

**Figure 3.**
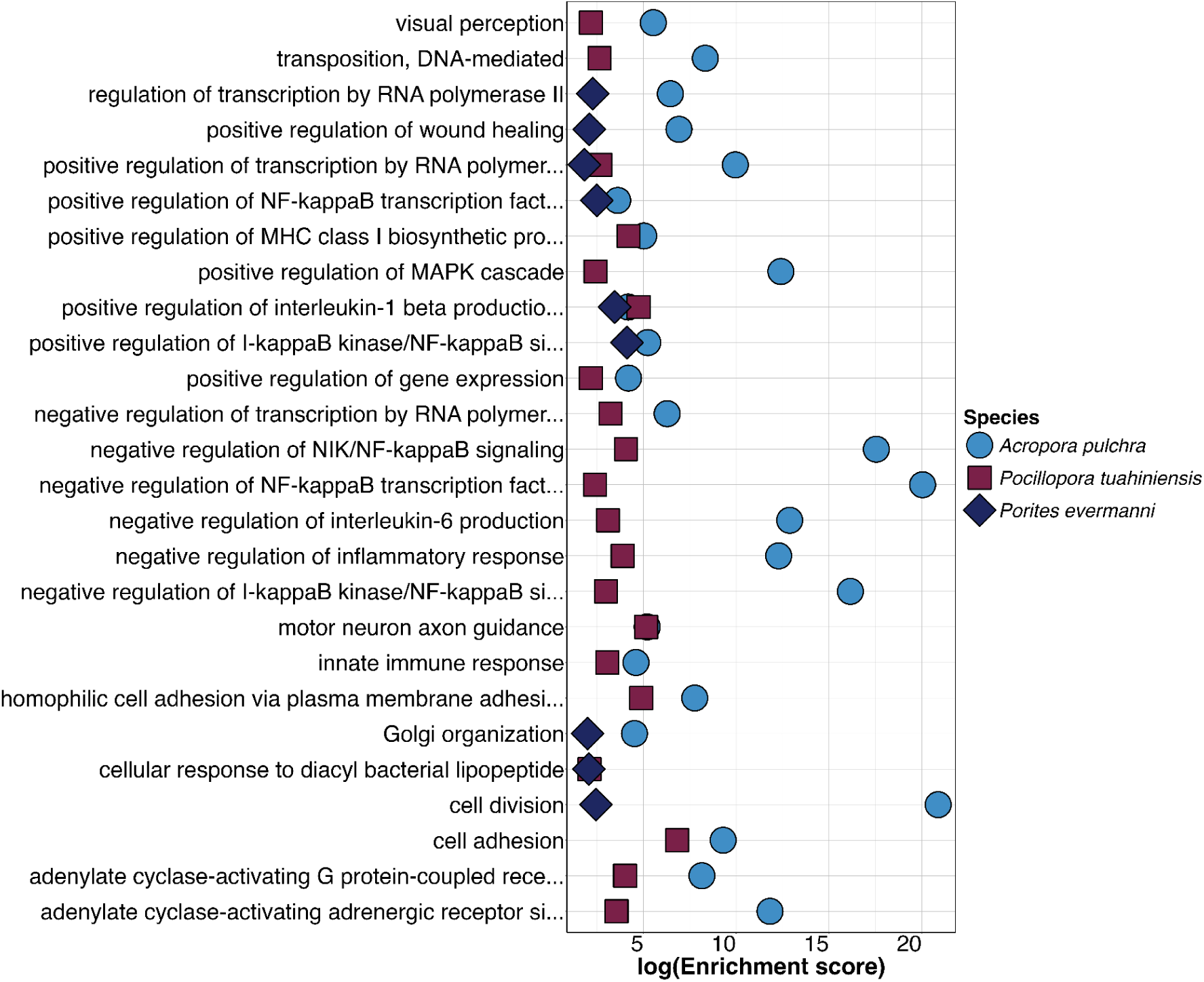
Enriched GO terms (biological processes) in genes closest to lncRNAs in three coral species.

**Table 2.** Percentages of ncRNAs that overlap (≥1bp overlap) with a gene.

|  | *A. pulchra* | *P. evermanni* | *P. tuahiniensis* |
| --- | --- | --- | --- |
| lncRNA | 1.1% | 2% | 4.3% |
| piRNA | 71.3% | 84.7% | 99.2% |

In all species, a majority of piRNAs clusters (85.1%) overlapped with genes (Table 2). In all species, genes that overlapped with piRNA clusters were enriched for GO biological processes related to DNA functions. However, no specific enriched GO terms were enriched across all species. For genes closest to piRNA clusters, *A. pulchra* and *P. tuahiniensis* shared enriched GO biological processes, including regulation of mitochondrial and DNA-templated transcription, DNA recombination, and DNA integration (Figure 4; Table S5). In contrast, *A. pulchra* and *P. evermanni* shared enriched GO processes related to telomere maintenance, G-quadruplex DNA unwinding, and DNA unwinding during DNA replication (Figure 4; Table S5). No GO biological processes were found to be shared between *P. evermanni* and *P. tuahiniensis*.

**Figure 4.**
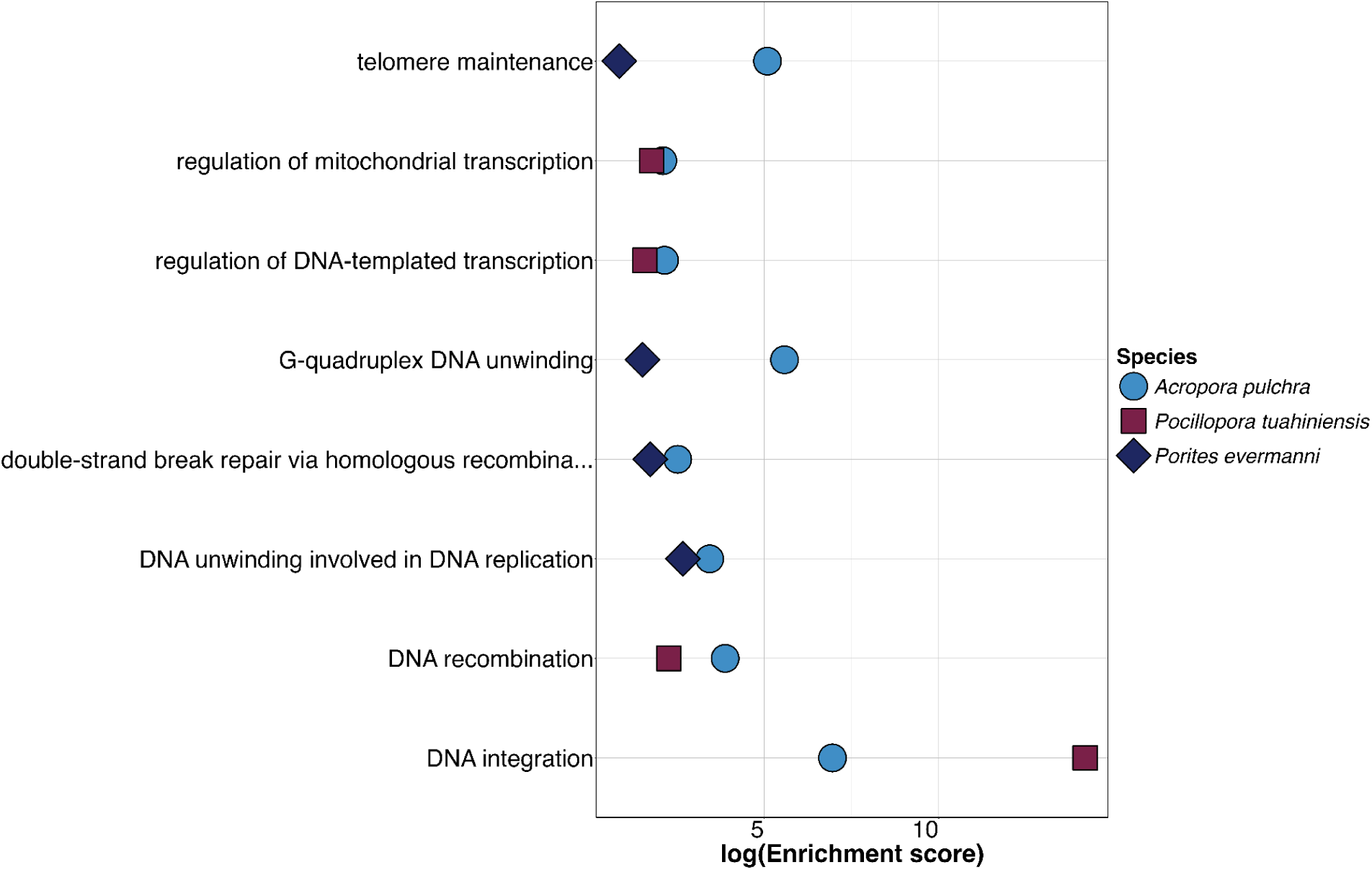
Enriched GO terms (biological processes) of the genes overlapping with piRNAs clusters in three coral species.

### miRNA target identification

To identify processes that are potentially regulated by miRNAs in the three species, a GO enrichment analysis was performed on the predicted gene targets of the miRNAs in each species. *A. pulchra* and *P. tuahiniensis* had the highest number of shared enriched GO biological processes (6 terms) including response to bacterium, negative regulation of type I interferon-mediated signaling pathway, motor neuron migration, generation of precursor metabolites and energy, bioluminescence, and axonal fasciculation (Figure 5; Table S6). *A. pulchra* and *P. evermanni* shared two GO biological processes, homophilic cell adhesion via plasma membrane adhesion molecules and defense response to virus (Figure 5; Table S6). *P. evermanni* and *P. tuahiniensis* shared positive regulation of MAPK cascade and cAMP transport GO terms (Figure 5; Table S6).

**Figure 5.**
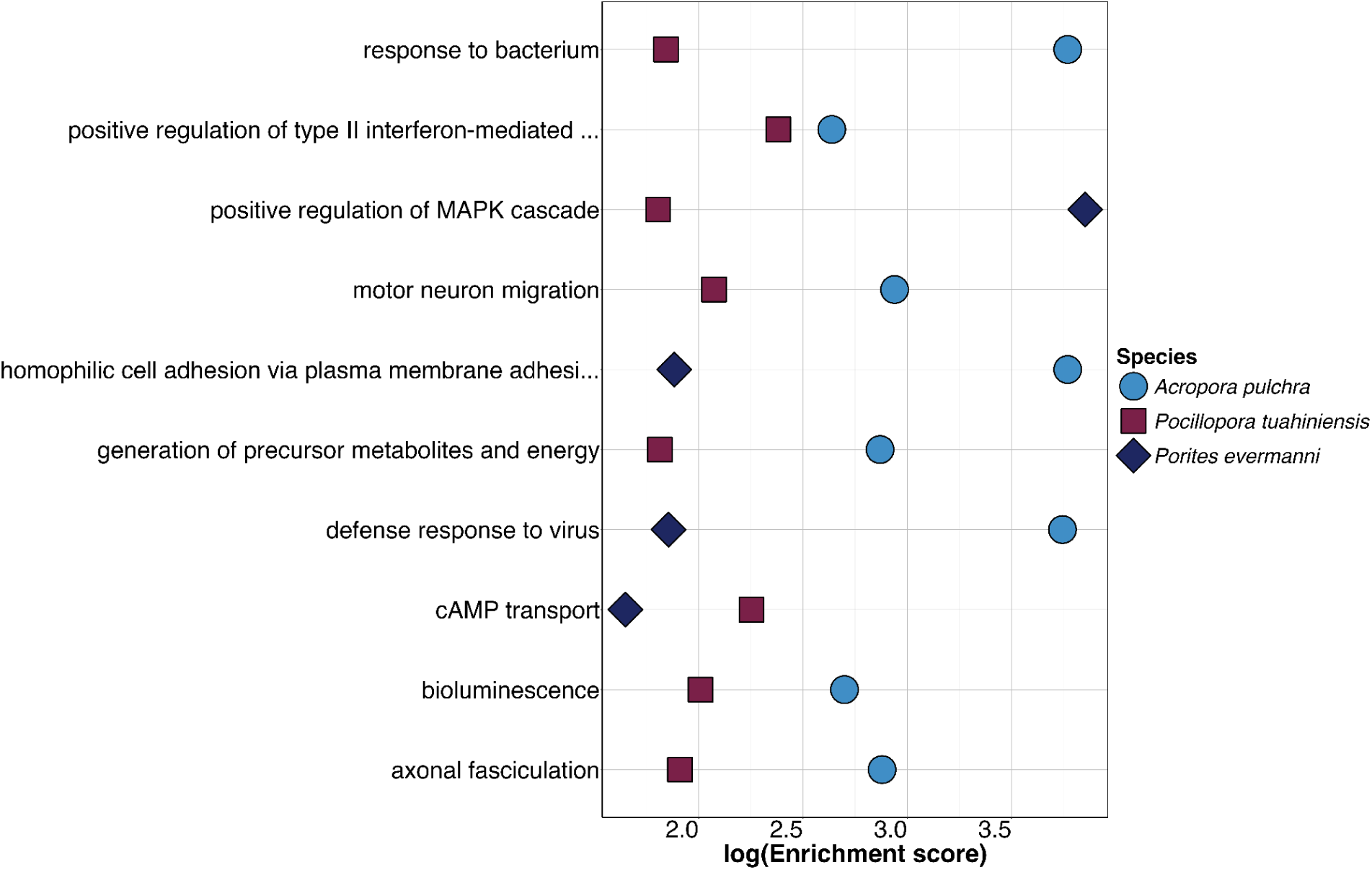
Enriched GO terms (biological processes) of the genes targeted by miRNAs in three coral species.

GO enrichment was also evaluated for the miRNAs shared across the three species: mir-100, mir-2023, mir-2025, and mir-2036. Putative targets of mir-100 were enriched for vitamin D receptor signaling pathway, regulation of presynaptic cytosolic calcium ion concentration, and acetyl-CoA biosynthetic processes, in *A. pulchra* and *P. evermanni* (Figure 6; Table S7). The term sporulation resulting in formation of a cellular spore was enriched in genes targeted by mir-100 in *P. evermanni* and *P. tuahiniensis* (Figure 6; Table S7). Only *P. evermanni* and *P. tuahiniensis* shared enriched GO terms in the genes targeted by mir-2023; the GO terms negative regulation of T cell mediated immune response to tumor cell, cellular response to forskolin, and cellular response to 2,3,7,8-tetrachlorodibenzodioxine were enriched (Figure 6; Table S7). For *P. evermanni* and *P. tuahiniensis*, genes targeted by mir-2025 were enriched in processes related to positive regulation of transcription by RNA polymerase II and negative regulation of urine, while mir-2025 targeted genes enriched in coronary artery morphogenesis, and cholesterol catabolic process in *A. pulchra* and *P. tuahiniensis* (Figure 6; Table S7). Finally, targets of mir-2036 were enriched in processes including positive regulation of insulin secretion for *A. pulchra* and *P. tuahiniensis*, and homophilic cell adhesion via plasma membrane adhesion molecules for *P. tuahiniensis* and *P. evermanni*.

**Figure 6.**
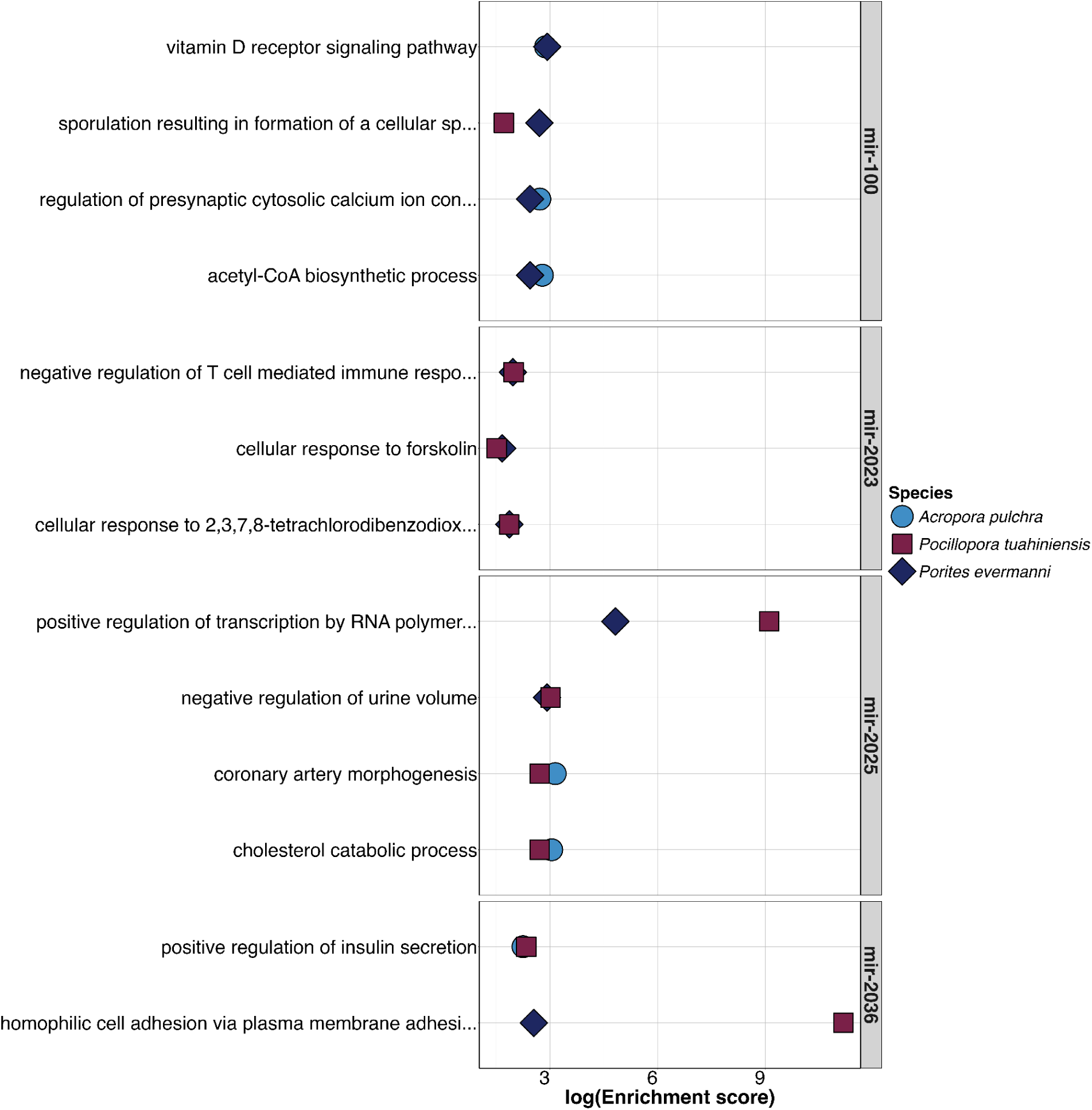
Enriched GO terms (biological processes) of the genes targeted by conserved miRNAs (mir-100, mir-2023, mir-2025, mir-2036) in three coral species.

## Discussion

The characterization of a diverse ncRNAs repertoire in *A. pulchra*, *P. evermanni*, and *P. tuahiniensis* provides a novel perspective on gene expression regulation via ncRNAs and its role in coral acclimatization, which influences the ability of different coral species to survive in a variety of ecological conditions. The identification of conserved ncRNA machinery components in the *A. pulchra*, *P. evermanni*, and *P. tuahiniensis* annotated proteomes indicates the conservation and functional importance of sophisticated ncRNA-mediated gene regulatory mechanisms. Few lncRNA and miRNA sequences were shared across species, indiciating high evolutionary turnover for these ncRNAs, potentially contributing to the unique characteristics of each coral species. However, the shared enrichment of specific gene functions in multiple species highlights potential evolutionary conservation of regulatory interactions, even though the ncRNAs themselves do not have conserved sequences.

### ncRNA protein machinery present in study species

In all coral species in this study, critical miRNA biogenesis proteins, including Drosha, DCGR8, XPO5, and Dicer, were found (Table 1). The presence of these proteins confirm functional miRNA-processing protein machinery in corals, which support the production of functional mature miRNA products. AGO2, a key protein in the multiprotein RNA-induced silencing complex (RISC) utilized by miRNAs to bind complementary mRNAs for degradation or translation suppression (Höck & Meister, 2008), was also present in all coral species. Further, PIP5K1A, a protein that inhibits the transport of pre-miRNAs from the nucleus to the cytoplasm (Li et al., 2023), indicates a sophisticated level of post-transcriptional control of miRNAs. This complexity in coral gene regulation opens up new avenues for research into the evolution of these mechanisms across diverse taxa, potentially prompting investigations in other basal metazoans.

Piwi proteins, which are associated with piRNAs, were found in all species in this study. Mature piRNAs bind piwi proteins to form piRNA-induced silencing complexes (piRISCs) and silence transposable elements and other targets (Czech et al., 2018). While piRNAs were first described in only animal germ cells (Siomi et al., 2011), piRNAs and piwi proteins were more recently found to be present in both somatic and germ cells of the cnidarians *Hydra* and *Nematostella*, indicating putative stem cell properties and/or somatic functionality (Juliano et al., 2014; Praher et al., 2017; Teefy et al., 2020). Given that the RNA extractions in our study were performed on whole tissue samples, we were unable to assess differences in somatic and germ cells in *A. pulchra*, *P. evermanni*, and *P. tuahiniensis*, although germ cell abundance was likely low in March, given the spawning periods for these species (*A. pulchra* spawns September through November (Becker, 2024; Carroll et al., 2006); *P. evermanni* spawns December through February (Adjeroud et al., 2007); *P. tuahiniensis* spawns November through December (Harnay et al., 2023)). Nevertheless, prevalence of gene-derived piRNAs in our analysis (Figure 2B) suggest that piRNA function in corals is beyond the canonical function of regulating transposable elements in the germ cells. Further investigation is needed to understand the piRNA-piwi protein functions and dynamics in different coral tissues.

There are multiple mechanisms of lncRNA biogenesis, making it difficult to broadly identify conserved proteins associated with lncRNA biogenesis (Chao et al., 2022). Generally, the biogenesis of lncRNAs is comparable to that of mRNA, in that the lncRNA gets capped, polyadenylated, and spliced utilizing the same proteins as mRNAs (Y. Liu, Ding, et al., 2021). RNase P, while primarily recognized for cleaving the 5’ end of precursor tRNAs, can also form the mature 3’ end of lncRNAs by cleavage (Y. Liu, Ding, et al., 2021). RNase P was found in *A. pulchra*, *P. evermanni*, and *P. tuahiniensis*, indicating that non-canonical mechanisms of lncRNA biogenesis are present in corals (Y. Liu, Ding, et al., 2021). Identification of this protein indicates that mechanisms of lncRNA biogenesis are likely occurring in *A. pulchra*, *P. evermanni*, and *P. tuahiniensis*.

### Evolutionary insights and functional implications of coral lncRNAs

Our study identified comparable numbers of lncRNAs to other studies in cnidarians: 8,117 in *Acropora digitifera* (Huang et al., 2019), 11,206 in *Protolpalythoa variabilis* (Huang et al., 2017), 13,240 in *Palythoa caribaeorum* (Huang et al., 2017), 6,328 in *Montipora foliosa* (Y. Liu, Liao, et al., 2021), 904 in *Pocillopora damicornis* (Yu et al., 2021), 7,059 in *Pocillopora damicornis* (Guo et al., 2021), 16,206 in *Acropora pulchra* (this study), 7,378 in *Porites evermanni* (this study), and 12,395 in *Pocillopora tuahiniensis* (this study). Few lncRNAs sequences were shared (<2%) between *A. pulchra*, *P. evermanni*, and *P. tuahiniensis*, indicating that only a small portion of lncRNAs may be conserved across multiple coral species. This contrasts with previous work in the cnidarian order Zoantharia, in which over a third of lncRNAs found in *Palythoa caribaeorum* and *Protolpalythoa variabilis* were shared (Huang et al., 2017). The discrepancy between our results and previous findings may stem from the greater evolutionary distance between *A. pulchra*, *P. evermanni*, and *P. tuahiniensis*, compared to the relationship between *Palythoa caribaeorum* and *Protolpalythoa variabilis* (Reimer et al., 2006). These findings reveal significant evolutionary divergence in lncRNAs across diverse coral species and emphasize the importance of expanding comparative research on gene regulation mechanisms within cnidarians.

We identified 12,395 lncRNA transcripts in *P. tuahiniensis* using Illumina sequencing technology, a number notably higher than the lncRNAs reported for its congener *Pocillopora damicornis*: 904 using Nanopore sequencing (Yu et al., 2021) and 7,059 using PacBio sequencing (Guo et al., 2021). These differences may be a result of methodological factors, including extraction protocols, sequencing technologies, lncRNA identification workflows, and stringency of analyses (Mattick et al., 2023). However, the disparities may also reflect inherent biological differences between *P. tuahiniensis* and *P. damicornis*. Although they are members of the same genus, *P. tuahiniensis* and *P. damicornis* differ in morphology, physiology, and reproductive strategy (Johnston & Burgess, 2023; Schmidt-Roach et al., 2014), traits that could influence their lncRNA repertoires and their roles in gene expression regulation. Comparative analyses of lncRNAs across species provide critical insights into the evolutionary and ecological functions of these molecules, shedding light on how life history traits, environmental pressures, and genetic diversity shape lncRNA evolution. Addressing these knowledge gaps is essential to understanding the broader significance of lncRNAs in coral biology and their potential roles in adaptive responses to environmental change.

lncRNAs in *A. pulchra, P. evermanni,* and *P. tuahiniensis* are often in proximity to genes coding for immune-related proteins. This proximity suggests that lncRNAs may be involved in immune processes in the three species, as proximity to genes can often suggest functional relationships for lncRNA (Ma et al., 2013; Marchese et al., 2017). Immune functions in corals are known to play critical roles in bleaching susceptibility and disease resistance (Mansfield & Gilmore, 2019; Mydlarz et al., 2010; Weis, 2008). Previous studies have shown lncRNAs to be involved in critical coral biological processes such as thermal resilience (Yu et al., 2021), bleaching thresholds (Huang et al., 2017), and symbiont infection (Huang et al., 2019). For example, lncRNAs have been implicated in the post-transcriptional regulation of Ras signal transduction pathway proteins and components of the innate immune system in cnidarians (e.g., complement factor B and stimulator of interferon genes protein; (Huang et al., 2017)). The immune system, the primary role of which is to detect and regulate non-self organisms, is intrinsically linked to all stages of coral-Symbiodiniaceae relationships, including establishment, maintenance and breakdown (Davy et al., 2012; Helgoe et al., 2024; Weis, 2008). Given this connection, lncRNA regulation of immune-related genes suggests that lncRNAs may influence symbiosis, potentially affecting symbiont recognition, stability or expulsion during bleaching (Huang et al., 2017). Alternatively or in parallel, lncRNAs could be involved in regulating immune genes related to disease response, another major threat to coral health (Mydlarz et al., 2010; Sweet & Bulling, 2017). Thus, lncRNAs may serve as key regulators of coral resilience by influencing symbiosis and immune defense mechanisms in response to environmental stressors.

### Few miRNAs are conserved across species, but share enriched functions of targets

Four miRNAs (mir-100, mir-2023, mir-2025, mir-2036) were conserved across all three species of interest and corresponded to miRNAs that have been identified in other cnidarians (Praher et al., 2021). mir-100 has been identified in many taxa, including bilaterians and cnidarians (Grimson et al., 2008; Praher et al., 2021). In our analyses, genes targeted by mir-100 were enriched in vitamin D receptor signaling pathway, regulation of presynaptic cytosolic calcium ion concentration, and acetyl-CoA biosynthetic process in *A. pulchra* and *P. evermanni*, as well as sporulation resulting in formation of a cellular spore in *P. evermanni* and *P. tuahiniensis*. These enriched processes demonstrate that while mir-100 may be highly conserved, the gene targets of mir-100 can differ by species, highlighting the diverse regulatory roles that mir-100 likely plays in corals. Previous studies have highlighted the versatility of mir-100 in cnidarians. In *Stylophora pistillata*, mir-100 targeted genes homologous to those related to embryonic forelimb morphogenesis and bone development, which the authors hypothesized importance in coral calcification (Liew et al., 2014). Additionally, mir-100 has been shown to be expressed in the oral end of *Nematostella* larvae, demonstrating tissue-specific expression patterns during development (Moran et al., 2014). In both adult and larval *Acropora digitifera*, mir-100 is expressed (Gajigan & Conaco, 2017). Although Gajigan and Conaco 2017 did not predict specific targets of mir-100 in their study, the presence of mir-100 across life stages implies diverse and potentially distinct roles throughout coral development. Together, these findings underscore the evolutionary conservation of mir-100 in corals while also revealing its diverse regulatory roles in multiple coral species.

When examining all miRNAs and their putative targets, *A. pulchra* and *P. tuahiniensis* demonstrated the greatest overlap in the biological processes associated with their target genes. In both species, miRNAs targeted genes with enriched GO terms such as response to bacterium and negative regulation of type I interferon-mediated signaling pathway, suggesting that miRNAs play a role in immune response regulation and immune signaling pathways. In model organisms, miRNAs are increasingly viewed as critical regulators of innate and adaptive immune responses, and their abnormal expression has been linked to human diseases and cancers (Raisch et al., 2013). In *P. evermanni* and *P. tuahiniensis*, gene targets of miRNAs shared enriched GO terms such as regulation of MAPK cascade and cAMP transport, indicating potential miRNA roles in signal transduction pathways and cellular communication. The identification of diverse yet overlapping GO terms across species indicates that miRNAs may be finely tuned to species-specific needs while still preserving certain essential functions. This functional diversity could reflect the adaptive role of miRNAs in modulating responses to species-specific environmental challenges, which might include differences in habitat, symbiotic relationships (Baumgarten et al., 2013, 2018), or life history strategies.

### Characteristic piRNAs biogenesis and possible regulation of genome maintenance and stability

The putative piRNAs identified in *A. pulchra*, *P. tuahiniensis*, and *P. evermanni* had characteristics indicative of piRNA ping-pong biogenesis, including an adenine in the 10th position in the piRNA transcripts and the presence of a uracil on the 5’ end of piRNA transcripts (Czech & Hannon, 2016). The piRNA ping-pong mechanism involves an amplification loop in which primary piRNAs, bound to piwi proteins, recognize and cleave target transposon RNAs, generating secondary piRNAs with complementary sequences (Czech et al., 2018). The cleavage typically occurs 10 bp downstream of the 5’ end of the primary piRNA, resulting in an adenine in the 10th position (Czech et al., 2018). The presence of a 5’ uracil is linked to the binding specificity of the piwi proteins, which preferentially bind to piRNAs with a 5’ uracil, leading to easier loading and stabilization of piRNAs in the piRISC structure (Stein et al., 2019). This cycle not only amplifies piRNAs but also provides a robust defense against transposable elements and maintains genomic stability (Teefy et al., 2020). The conservation of these signatures across coral species suggests a fundamental role for piRNAs in genome defense.

No enriched GO terms were shared across all species in genes closest to piRNA clusters. However, genes closest to piRNA clusters in *A. pulchra* and *P. tuahiniensis* shared enriched GO terms, including regulation of mitochondrial and DNA-templated transcription, DNA recombination, and DNA integration. Additionally, enriched functions in genes closest to piRNA clusters in *A. pulchra* and *P. evermanni* included telomere maintenance, G-quadruplex DNA unwinding, and DNA unwinding involved in DNA replication. piRNAs bind to piwi proteins to either silence transposons or genes, indicating potential piRNA-dependent gene regulations (Simonelig, 2014). Thus, piRNAs in *A. pulchra*, *P. tuahiniensis*, and *P. evermanni* may be important in regulating genes related to genomic stability, as seen in other taxa (Moyano & Stefani, 2015; Van Wolfswinkel & Ketting, 2010). The enrichment of shared terms such as DNA recombination and integration in *A. pulchra* and *P. tuahiniensis* and telomere maintenance and G-quadruplex DNA unwinding in *A. pulchra* and *P. evermanni* suggests that there is a conserved role for piRNAs in maintaining genomic stability and integrity across species (Czech et al., 2018). The overlap of piRNAs with genes associated with transcription, DNA maintenance, and genomic stability highlights the potential for piRNA-mediated regulation of these essential processes, potentially safeguarding the genomes of diverse coral species.

### ncRNAs as epigenetic regulators

ncRNAs are key players in epigenetic regulation, which encompasses molecules and mechanisms that can perpetuate different gene activity states without a change in the DNA sequence (Cavalli & Heard, 2019). ncRNAs can modulate the expression of key epigenetic actors, including DNA methyltransferases, histone deacetylases, histone methyltransferases and others (Bure et al., 2022; Joh et al., 2014; Sadida et al., 2024; Xie et al., 2023; Yao et al., 2019). The expression of ncRNAs themselves are also subject to regulation by DNA methylation, RNA modification and histone modifications (Bure et al., 2022; Peschansky & Wahlestedt, 2014; Sadida et al., 2024; Xie et al., 2023; Yao et al., 2019). These reciprocal regulatory relationships mediate critical cellular functions and exert extensive influence on gene expression, ultimately shaping phenotypic variation among species and individuals (Mattick and Makunin 2006). Future research should prioritize investigating interactions between ncRNAs and other epigenetic regulators in corals, as these mechanisms may play a critical role in acclimatization and adaptation to climate change.

Transgenerational epigenetic inheritance occurs when epigenetic features are transmitted via the germline from one generation of organisms to at least two generations of offspring (Bošković & Rando, 2018) and has been proposed as a possible mechanism for rapid coral acclimatization to climate change (Putnam, 2021; Torda et al., 2017). While RNAs themselves do not necessarily act as the heritable epigenetic signal, they are crucial in the establishment of inheritable DNA methylation or histone patterns (Fitz-James & Cavalli, 2022). For example, in fission yeast, multiple lncRNAs compete with H3K9me for binding to Swi6, a protein involved in heterochromatic gene silencing during mitosis and meiosis (Motamedi et al., 2008). By preventing the spreading of Swi6 and subsequent K3K9 methylation, lncRNAs directly influence chromatin organization across generations (Keller et al., 2012). Similarly, in *C. elegans*, piRNAs dictate highly stable long-term gene silencing that can persist for at least 20 generations (Ashe et al., 2012). ncRNAs can be transmitted to the next generation both maternally through eggs (Brennecke et al., 2008; Roovers et al., 2015; Tang et al., 2007) and paternally through sperm (Conine et al., 2018; X. Zhang et al., 2017). This suggests that ncRNAs may serve as critical secondary signals from which other epigenetic signals might be reconstructed (Fitz-James & Cavalli, 2022).

Intriguingly, evidence suggests that ncRNAs can be exchanged across species via extracellular conveyors, a phenomenon that may be particularly relevant for reef-building corals due to their complex holobiont structure (Di Liegro et al., 2017; Leitão et al., 2020). For instance, in response to infection by *Verticillium dahliae*, a fungal pathogen, plants upregulate miR-166 and miR-159 and export these to the fungus, where they silence genes critical for fungal virulence, including a Ca2+-dependent cysteine protease and an isotrichodermin C-15 hydroxylase (T. Zhang et al., 2016). In corals, similar cross-kingdom ncRNA communication could play a role in epigenetic regulation in the holobiont, influencing interactions between the coral host, symbionts, and associated microbes (Leitão et al., 2020). Previous studies have demonstrated differential expression of miRNAs and lncRNAs in response to symbiotic infection in the sea anemone *Aiptasia* and *Acropora digitifera*, respectively (Baumgarten et al., 2018; Huang et al., 2019). This suggests that ncRNAs may actively mediate symbiotic relationships or respond to them, potentially facilitating cross-kingdom communication and contributing to the regulatory complexity of the coral holobiont.

Although epigenetic crosstalk remains largely unexplored in corals, it holds promising potential as a mechanism of plasticity for corals to respond to environmental perturbations. Preliminary evidence suggests that environmentally mediated epigenetic crosstalk may occur in at least one coral species. In response to transplantation, *Porites astreoides* exhibited differentially methylated loci associated with ncRNA sequences in the genome, suggesting potential regulatory relationships between DNA methylation and ncRNAs under environmental change (Dimond & Roberts, 2020). By identifying and characterizing ncRNAs in the three coral species examined here, we provide a foundational resource for investigating relationships between epigenetic regulators.

### Limitations of analyses

While we provide a comprehensive overview of coral ncRNAs, we acknowledge the limitations of the analyses presented. Our study species presented varying degrees of genomic quality and completeness, which may have contributed to differences observed among species. For example, the *A. millepora* genome, which was used for the *A. pulchra* analysis in this study, has more genomic elements annotated than the other genomes used in this study. Additionally, there was discrepancy in the proportion of known miRNAs for each species. Using the cnidarian database and miRBase, *A. pulchra* had more known miRNAs (24 known miRNAs; 68% of total miRNAs identified) than *P. evermanni* (nine known miRNAs; 19% of total miRNAs identified) or *P. tuahiniensis* (nine known miRNAs; 24% of total miRNAs identified). This higher rate of database matching in *A. pulchra* is likely related to the composition of the reference database, rather than a true difference in miRNA conservation. Our curated reference of known cnidarian miRNAs contained 90 *Acropora* spp. miRNAs, but zero known miRNAs identified in *Porites* or *Pocillopora*. Further, miRBase does not contain known sequences from any of these three study genera.

Methods to characterize gene expression in corals are well established, but standardized methodologies and tools to describe gene expression regulation in corals are still in their early stages. Most tools for ncRNA identification or analysis are designed for model systems, which may rely on assumptions that may not apply to ncRNA dynamics in corals. Additionally, the choice of identification method or tool can affect the number of valid ncRNAs identified, as different methodologies often apply varying filters and classification thresholds. Here, we applied stringent filters across all tools to ensure greater confidence in the ncRNA candidates identified and to provide a robust framework for further analysis in coral systems.

## Conclusion

Our comprehensive characterization of the ncRNA repertoire in three diverse coral species provides valuable insights into the regulatory landscape of these ecologically crucial organisms. The identification of conserved ncRNA protein machinery and specific ncRNA sequences across species suggests a fundamental role for these regulatory elements in coral biology. The functional enrichment of genes associated with lncRNAs, piRNAs, and miRNA targets points to potential involvement in key processes such as immunity, DNA maintenance, transcription, and metabolism. Not only do these findings expand on our current understanding of coral molecular biology, but they also open new avenues for investigating the epigenetic mechanisms underlying coral acclimatization and resilience to environmental stressors. Future research should focus on elucidating specific functions of ncRNAs and their potential roles in mediating responses to climate change-induced stressors and pathogens.

## Acknowledgements

This work was supported by the National Science Foundation (NSF) Rules of Life-Epigenetics Awards to HMP (1921465), JEL (1921402), and SBR (1921149), as well as NSF Graduate Research Fellowships to JA and DMB, and supported by resources from NSF-OCE 2224354 to the Mo’orea Coral Reef LTER, as well as a generous gift from the Gordon and Betty Moore Foundation. We thank Kristina Terpis for laboratory assistance. The authors acknowledge use of the resources of the URI Center for Computational Research and the FIU Instructional and Research Computing Center for this work. As guests, we honor and acknowledge the Mā’ohi community, we give thanks for the land and water resources of Polynesia and the traditional custodians of the land on which this experimental work was conducted on the island of Mo’orea. Māuruuru roa. With respect to the spelling of Tahitian words, we endeavored to followed the Te Fare Vānaʻa, transcription system that is adhered to by a large segment of the Tahitian community, but also recognize other community members follow the Raapoto transcription system where the island name of Moorea is, for example, spelled without the ‘eta (i.e., Moorea).

## Data availability statement

Data and scripts are available on Github at https://github.com/urol-e5/deep-dive. Sanger sequences and archived genomic references have been deposited at https://osf.io/aw53f/. RNA sequences and small RNA sequences are stored on NCBI under BioProjects PRJNA1236658 and PRJNA1236666, respectively.

