## Supplemental Methods & Figures for "Non-coding RNA Repertoire in Reef-Building Corals"

While *Acropora pulchra* can be reliably identified by morphology and we have confirmed fertilization success of *A. pulchra* from the collection site (Becker, 2024), *Pocillopora spp.* and *Porites spp.* are known to include cryptic species, requiring genetic identification (Forsman et al., 2009; Burgess et al., 2021). *Pocillopora* species were identified by amplifying the mitochondrial open reading frame (mtORF) region as described by (Johnston et al., 2018; Burgess et al., 2021) using primers from (Flot et al., 2008): FatP6.1 (5′-TTTGGGSATTCGTTTAGCAG-3′) and RORF (5′-SCCAATATGTTAAACASCATGTCA-3′). Master mixes contained 12.55 µL of EmeraldAmp GT PCR Master Mix (TaKaRa Bio USA Inc. Cat # RR310B), 0.32 µL of forward and reverse primers as listed, 1 µL of template DNA, and 10.80 µL of nuclease free water (Invitrogen UltraPure CAT # 10977015) totaling 25 µL for the final volume. Positive controls were included as previously successively amplified gDNA samples using the primers listed and negative controls were included as master mix without template DNA. mtORF was amplified using a polymerase chain reaction (PCR) profile of a single denaturation set of 94°C for 60 seconds followed by 30 cycles of 94°C for 30 seconds for denaturation, 53°C for 30 seconds for annealing, and 72°C for 75 seconds for extension and a final incubation of 72°C for 5 min. PCR products were assessed with a 1.5% agarose gel in TAE for 30 minutes at 80 volts.

*Porites* species were identified using the coral nuclear histone region spanning H2A to H4 (i.e., H2; (Tisthammer et al., 2020)) using the following primers: zH2AH4f (5′-GTGTACTTGGCTGCYGTRCT-3′) and zH4Fr (5′-GACAACCGAGAATGTCCGGT-3′). The coral nuclear histone regionH2 was amplified using a PCR profile of a single denaturation set of 94°C for 2 minutes followed by 34 cycles of 96°C for 20 seconds for denaturation, 58.5°C for 20 seconds for annealing, and 72°C for 90 seconds for extension and a final incubation of 72°C for 5 minutes. Products were assessed on a gel as described above with an expected band size of approximately 1500 bp. DNA was assessed with a 1.5% agarose gel in TAE for 30 mins at 80 volts to confirm only one band of approximately 1500 bp was recovered.

One *Acropora pulchra* sample was sequenced for molecular markers previously used to classify *Acropora* species including the *Pax*-C 46/47 intron in the nuclear genome region as described by XXX as well as the mitochondrial putative control region (933+ bp) plus 83 bp of cytochrome oxidase III as described by XXX. The *Pax*-C intron region was amplified using the following primers: PaxC_intron-FP1 (5’-TCCAGAGCAGTTAGAGATGCTGG-3’) and PaxC_intron-RP1 (5′-GGCGATTTGAGAACCAAACCTGTA-3′; XXX). The mitochondrial control region was amplified using the following primers: CRf (5′-GCTTAGACAGGTTGGTTGATTGCCC-3′) and CO3r (5′-CTCCCAAATACATAATTTGAACTAA-3′; XXX). The PCR protocol for both regions was as follows: a single denaturation set of 95°C for 3 minutes followed by 35 cycles of 94°C for 30 seconds for denaturation, 53°C for 30 seconds for annealing, and 72°C for 60 seconds for extension and a final incubation of 72°C for 5 minutes. DNA was assessed with a 1.5% agarose gel in TAE for 30 mins at 80 volts. *Pax*-C amplification is variable in length (xxx) and resulted in bands of either a short (approx. 600 bp) or long (approx. 800 bp) length while the mitochondrial putative control region resulted in a single band approx. 1016 bp in length.

PCR products for all species were cleaned using ethanol precipitation, with 1/10th of the volume of 3M sodium acetate (Fisher Cat. AAJ61928AE) added to each PCR product, followed by an addition of 3 times the total volume of the mixture of ice-cold 100% ethanol. The mixture was incubated overnight at -20ºC and DNA was precipitated by centrifugation at 15,000 rcf for 30 minutes at room temperature. The pellet was washed with 70% ethanol twice, dried, and resuspended in 30 µl of 1M Tris-HCI, pH 8.0 (Fisher Cat. 15568025). Sanger sequencing using the same primers utilized during PCR amplification was performed at the URI Genomics and Sequencing center using Applied Biosystems BigDye Terminator v3.1. Sequences were aligned and analyzed using Geneious Alignment in GENEIOUS PRIME 2020.2.4. Pairwise alignment of reads was completed using Clustal Omega (Sievers et al., 2011) and neighbor-joining trees were constructed (Zhang & Sun, 2008) with Jukes-Cantor genetic distance model (Jukes & Cantor, 1969) for both forward and reverse reads for *Porites spp*.

**Supplemental Figures**

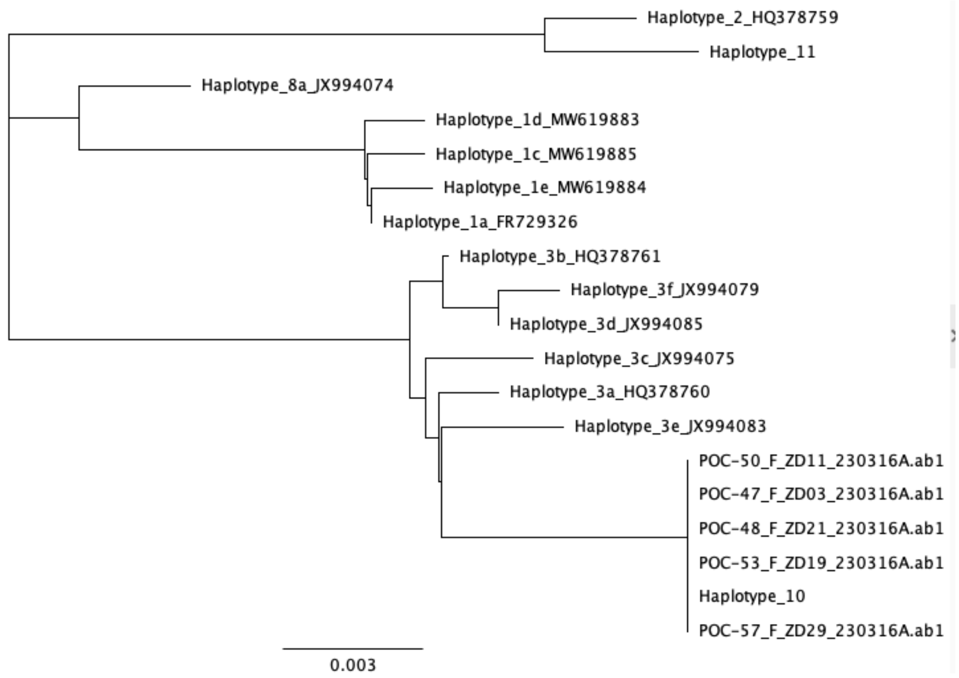


**Figure S1**. Species tree of the genetic relatedness of mtORF sequences was used to identify all of the samples as *Pocillopora tuahiniensis* (Haplotype 10; (Johnston & Burgess, 2023)). Samples in this project included those denoted by POC-#.

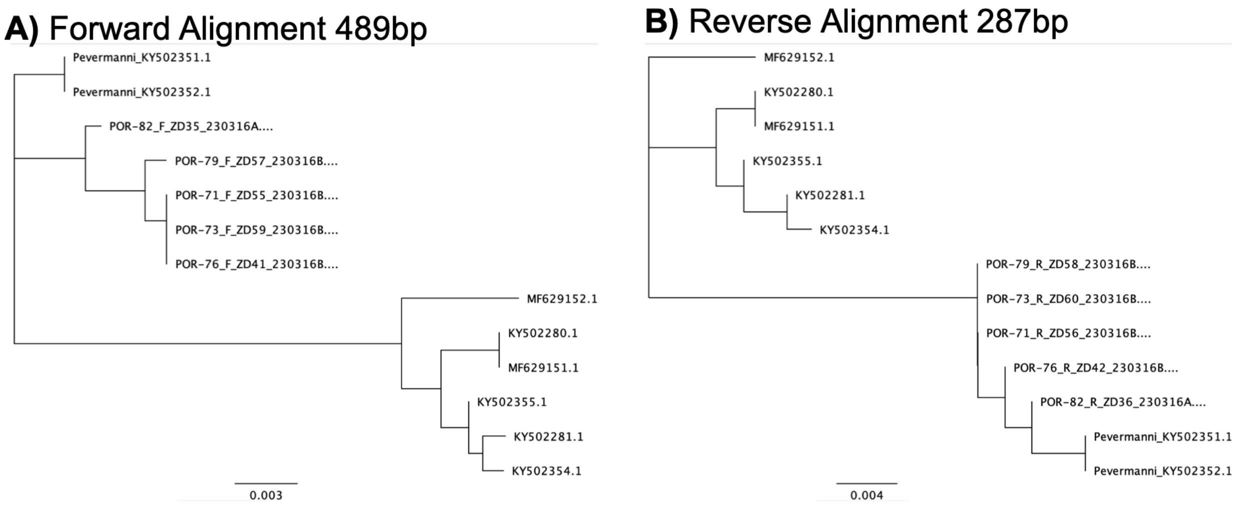


**Figure S2**. Coral nuclear histone region spanning H2A to H4 (H2) tree for broken down by **A)** forward reads and **B)** reverse reads. Samples in this project included those denoted by POR-#.

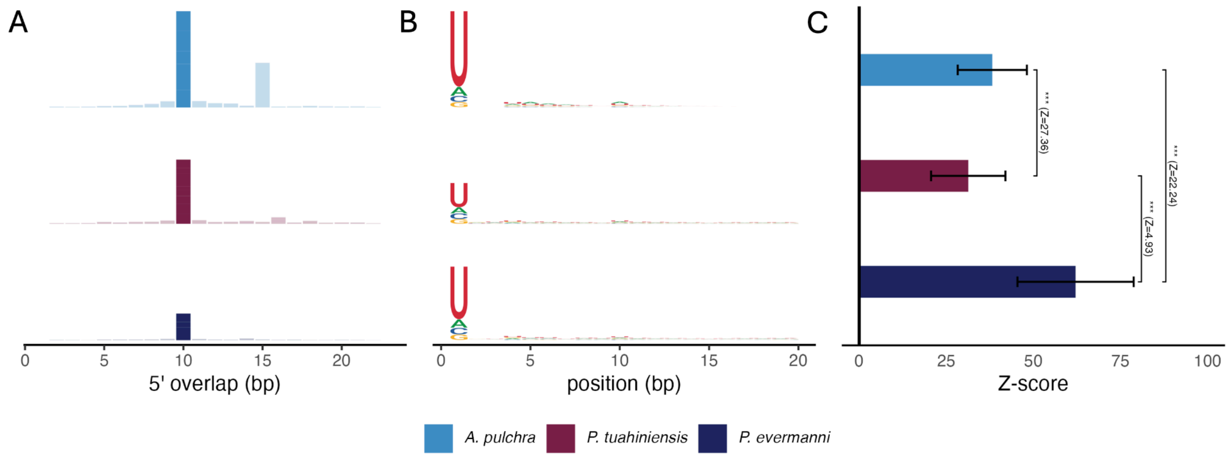


**Figure S3.** Putative piRNA characteristics in three coral species: *A. pulchra*, *P. tuahiniensis*, and *P. evermanni*. **A)** 5' overlap between complementary putative piRNA evidencing ping-pong secondary biogenesis. **B)** Base composition of putative piRNA showing 5'-end Uracil and position 10 Adenine biases typical of piRNAs**,** and **C)** Level of Ping-Pong amplification. Pairwise significance was assessed with a Mann-Whitney U-test.

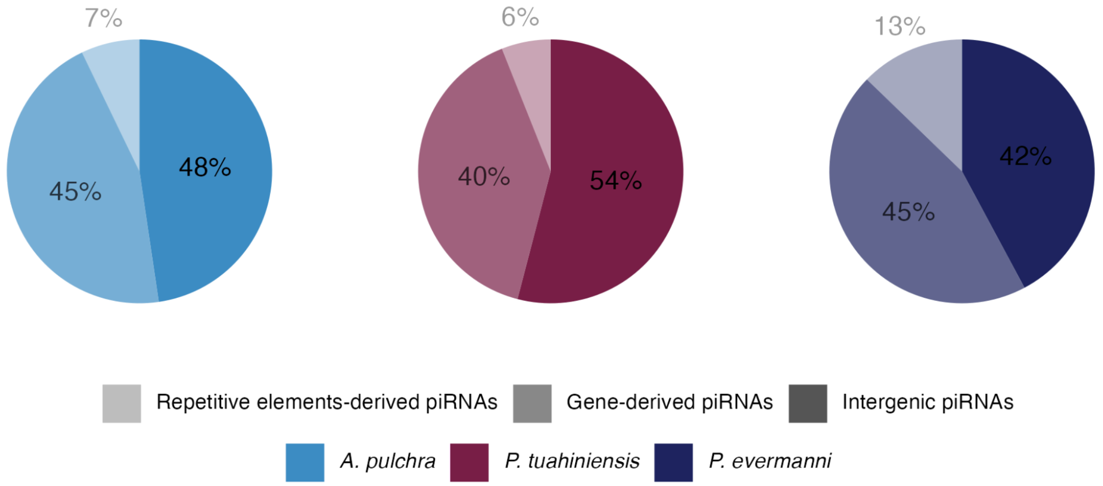


**Figure S4.** Proportions of gene-derived putative piRNA, repetitive element-derived putative piRNA, and intergenic putative piRNA reads in three coral species.
